# TomoTwin: Generalized 3D Localization of Macromolecules in Cryo-electron Tomograms with Structural Data Mining

**DOI:** 10.1101/2022.06.24.497279

**Authors:** Gavin Rice, Thorsten Wagner, Markus Stabrin, Stefan Raunser

**Affiliations:** Department of Structural Biochemistry, Max Planck Institute of Molecular Physiology, Otto-Hahn-Str. 11, 44227 Dortmund, Germany

**Author notes:** Both authors contributed equally (Gavin Rice, Thorsten Wagner).

## Abstract

Cryoelectron tomography enables the visualization of cellular environments in extreme detail through the lens of a benign observer; what remains lacking however are tools to analyze the full amount of information contained within these densely packed volumes. Detailed analysis of macromolecules through subtomogram averaging requires particles to first be localized within the tomogram volume, a task complicated by several factors including a low signal to noise ratio and crowding of the cellular space. Available methods for this task suffer either from being error prone or requiring manual annotation of training data. To assist in this crucial particle picking step, we present TomoTwin: a robust, first in class general picking model for cryo-electron tomograms based on deep metric learning. By embedding tomograms in an information-rich, high-dimensional space which separates macromolecules according to their 3-dimensional structure, TomoTwin allows users to identify proteins in tomograms *de novo* without manually creating training data or retraining the network each time a new protein is to be located. TomoTwin is open source and available at https://github.com/MPI-Dortmund/tomotwin-cryoet.

## Introduction

In recent years, cryo-electron tomography (cryo-ET) has emerged as a landmark technique for the visualization of macromolecules within their native cellular environment^1–7^. Advances in high-pressure freezing and the advent of focused ion beam (FIB) milling at cryogenic temperatures now allow for the routine preparation of thin (< 200 nm) lamellae from cells or even small organisms^8–10^. Performing cryo-ET on these thin lamellae offers a unique opportunity to capture cellular processes in 3D and in unprecedented detail. Subsequent analysis of specific macromolecules from tomographic volumes through subtomogram averaging (STA) allows in depth structural determination of macromolecular complexes in their native environment^11–14^. Particularly when complemented by recent advances in structure prediction such as alphafold2, STA forms a powerful crossbridge between protein biochemistry and cellular proteomics^15–17^. In order to perform STA however, particles of a macromolecule of interest must first be located within the tomographic volume, a task complicated by the 3D nature of these data.

The accurate localization of macromolecules inside cryo-electron tomograms is a well-recognized barrier for studying cellular life at the mesoscopic level, sparking competitions such as the annual Classification in Cryo-Electron Tomograms (SHREC) competition where contestants submit algorithms to localize proteins in tomograms with a benchmark set by template matching^18^. This has led to the development of several deep learning-based tools with high picking accuracies often achieved by leveraging popular 3D-Unet convolutional neural network (CNN) architectures^19–21^. Each of these approaches is unified however in the fact that they share a non-generalizing workflow, meaning that for each protein of interest, users must first manually pick the protein in at least one tomogram and retrain the neural network to identify that protein. Not only is this incompatible with the future directions of automated tomogram reconstruction and STA, but for many proteins picking sufficient training data by eye is not possible. With a minimal requirement for user-input, template matching^22, 23^ is still often utilized in cryo-ET processing workflows that place an emphasis on throughput^24^ although at the cost of picking accuracy.

One method to retain the accuracy of deep learning-based picking while circumventing the requirement of manually annotating training data for each protein of interest is to train a model to learn a generalized representation of 3D molecular shape that then can differentiate between macromolecules based on their structure. Such approaches have demonstrated profound impact for particle picking in 2D for single particle cryo-electron microscopy analysis^25–28^.

Particularly well suited for this type of generalization is deep metric learning in which data are encoded as a high-dimensional representation, called an embedding, where one or more learned characteristics of the data are related to distance in the embedding space^29, 30^. During training, the model is penalized for placing data from different classes near to one another and rewarded for placing data from the same class close together in the embedding space^31^. Therefore, over the training process the model learns to place data from each class within a distinct region of the embedding space where more similar classes are placed closer together and dissimilar ones further apart. In some cases, the embeddings of a dataset are sufficiently ordered to allow for *de novo* identification of classes based on their clustering in the embedding space^31^. By understanding similarity relationships, deep metric learning models have demonstrated acute adaptability when presented with new classes of data, being able to place them in the embedding space according to their similarity to known classes without requiring retraining^31–33^.

Here we present TomoTwin, a generalized particle picking model and deep metric learning toolkit for structural data mining of cryo-electron tomograms. We supply two workflows for macromolecular localization with TomoTwin, a reference-based workflow in which a single molecule is picked for each protein of interest and used as a target, and a *de novo* clustering workflow where macromolecular structures of interest are identified on a 2D manifold. Trained on a diverse set of simulated tomograms, the picking model of TomoTwin is able to locate new proteins with high accuracy in not only simulated data, but experimental and cellular tomograms as well. TomoTwin combines the high accuracy of deep learning-based particle picking with high throughput processing by removing the step of manual annotation of training data and model training, and allows simultaneous picking of several proteins of interest in each tomogram.

## Results

### Overview of functions, build, and philosophy behind TomoTwin

The machine learning backbone of TomoTwin is built on the principle of learning generalized representations of 3-dimensional shapes in tomograms (Supplementary Fig. 1a,b). Trained with deep metric learning, the 3D-CNN is able to locate not only macromolecules contained in the training set, but novel macromolecules in tomograms as well. This allows TomoTwin to retain the high fidelity of deep learning-based particle picking while avoiding the burden of requiring retraining for each protein of interest. The trained model plots tomogram subvolumes as points in a high-dimensional embedding space organized according to the similarity of their macromolecular contents (Supplementary Fig. 1c). Once this high-dimensional space is mapped for a tomogram, particles of each macromolecule can be picked by identifying their associated region in of the embedding space. This can be done either by identifying a single example of each protein of interest in a tomogram and using them to mark the region of the space where they are embedded to create a target embedding (reference-based workflow), or by plotting the tomogram embeddings onto a 2D manifold where clusters of subvolumes for each macromolecule can be identified by eye (clustering workflow) (Fig. 1a,b). Once the subvolumes containing a protein of interest are identified in the embedding space, they must be mapped back to real space in the tomogram where overlapping picks of the same molecule can be consolidated into one centralized pick per molecule (Fig. 1c). Finally, TomoTwin allows users to interactively filter the picked particles for each macromolecule of interest based on the particle size and the network’s confidence level, which is encoded as the distance between each subvolume and the target embedding for that macromolecule in the embedding space (Fig. 1d, Supplementary Fig. 2).

**Fig. 1:**
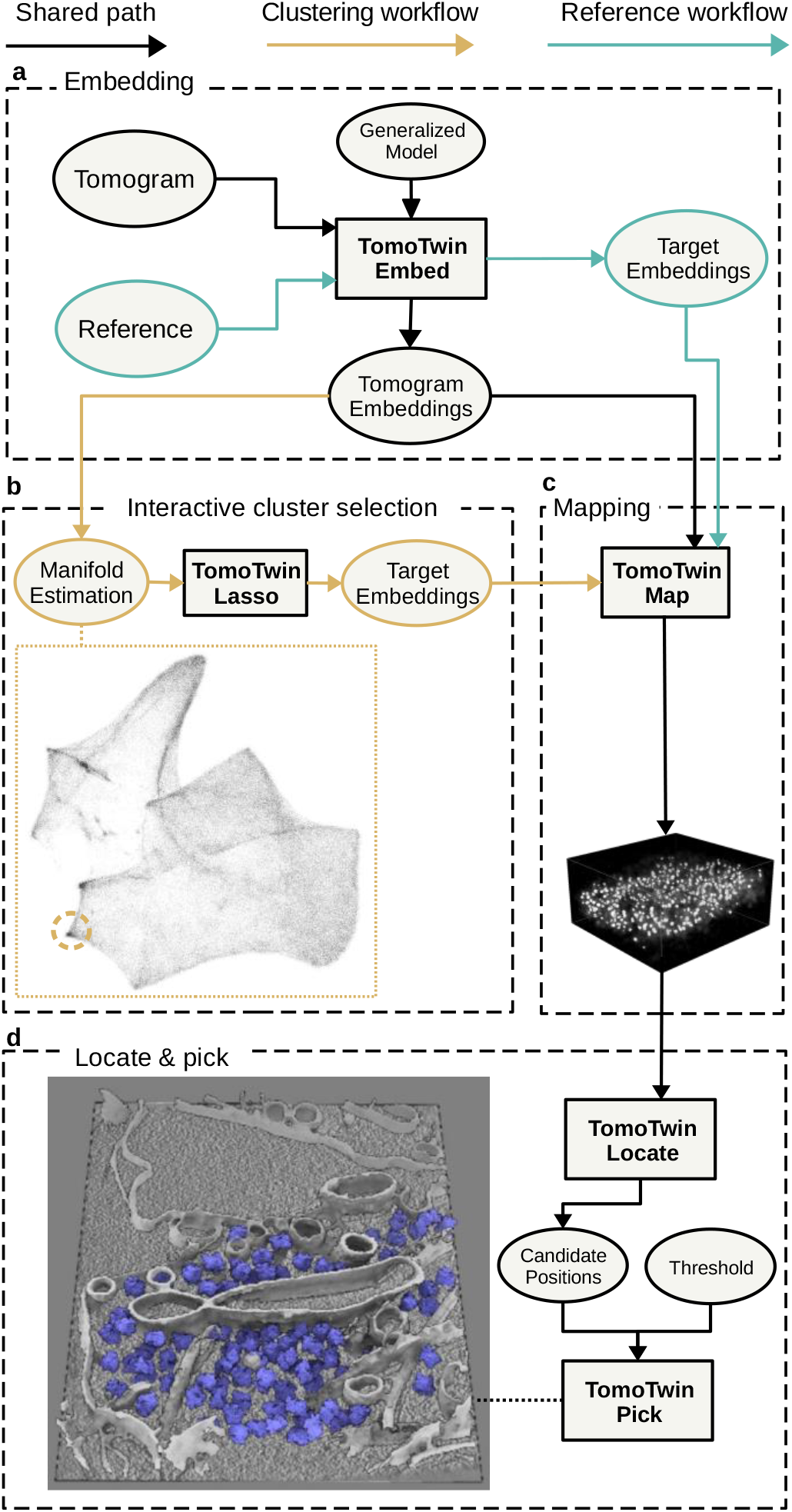
TomoTwin identifies and localizes particles by a clustering or a reference-based workflow. **a,** The first step in using TomoTwin is to embed the tomogram with the pre-trained model. Optionally, references can be selected and embedded as well to create target embeddings. **b,** For the clustering workflow the tomogram embeddings are projected on a 2D manifold and an interactive lasso tool is used to select clusters of interest to generate target embeddings. **c,** The distance matrix between each target embedding and the embeddings of the tomogram is calculated. **d,** All local maxima are located with TomoTwin Locate and are used to pick final coordinates for each protein of interest using TomoTwin Pick with confidence and size thresholding.

### Two workflows to identify and locate macromolecules in tomograms

TomoTwin represents tomograms in a high-dimensional space where subvolumes of each macromolecule are embedded in a distinct region of the space. In order to identify which region of the embedding space a macromolecule is located in, we provide the user with two workflows – a reference-based workflow and a clustering workflow. Each workflow picks particles with high accuracy, but the reference-based approach begins with identifying an example of the protein of interest in the tomogram and mapping this to the embedding space whereas the clustering workflow begins with identifying a region of the embedding space and mapping this to the tomogram. Which workflow is most suitable for any given application depends on how easily the protein(s) of interest can be identified in the tomogram versus the embeddings. Both workflows share the common first step of using the embedding function of TomoTwin to generate a high-dimensional embedding of the entire volume of the tomogram called a tomogram’s representation map (Fig. 1a).

In the reference-based workflow, users identify a single molecule of each protein of interest in a tomogram and embed it to generate a target in the embedding space for that protein. In the clustering workflow, TomoTwin approximates the representation map of the tomogram onto a 2-dimensional manifold. This 2D manifold can then be directly visualized by the user who can then outline one or more clusters of interest using the Lasso function of TomoTwin. The Lasso function then computes the center coordinate of the drawn cluster in the high-dimensional embedding space to be used as a target embedding in lieu of a reference (Fig. 1b). The map function of TomoTwin takes as input the tomogram embeddings and target embeddings and calculates the distance matrix between the target(s) and each point in the tomogram embeddings. The distances are mapped to the coordinates of each subvolume, constructing a similarity map of proposed particle locations within the tomogram for each protein of interest (Fig. 1c). The Locate function uses this similarity map to localize peaks of high similarity and generate candidate particle positions. Finally, the Pick function of TomoTwin uses these candidate positions as well as adjustable size and confidence thresholds to pick particles in the tomogram producing a coordinate file for each protein of interest to be then used for subtomogram averaging or other analysis (Fig. 1d).

### Training of the general picking model

To produce a picking model capable of localizing novel macromolecules within tomograms without requiring retraining, TomoTwin is trained using deep metric learning on triplets of subvolumes from simulated tomograms. A set of 120 structurally dissimilar proteins procured from the Protein Data Bank (PDB) ranging in size from 30 kDa to 2.7 mDa were used to simulate 84 tomograms containing a total of 120,000 subtomogram particles (Supplementary Fig. 3). During training, batches of subvolumes are embedded by a custom-built 3D CNN which transforms each 37×37×37 realspace 3D subvolume to a 1D, 32-length coordinate vector located on a high-dimensional embedding manifold molded to the surface of a 32D hypersphere (Supplementary Fig. 1a).

These coordinate vectors are used in the metric learning process which rewards the model for placing the anchor and positive close together in the embedding space and penalizes it for placing the anchor and negative near one another. Therefore, through training TomoTwin learns to place subvolumes of each macromolecule within a distinct region of the embedding space, where more structurally similar macromolecules are placed closer together and dissimilar ones further apart (Supplementary Fig. 1c). By training on a large, diverse set of 3D macromolecular shapes and sizes, TomoTwin learns a generalized representation of 3D macromolecular shapes which it leverages to place novel macromolecules in their own region of this embedding space relative to their structural similarity to known proteins without requiring retraining.

### The general picking model accurately locates particles across a wide range of shapes and sizes

Because *a priori* information on the ground truth locations of all molecules in a tomogram is not possible to obtain for experimental data, the picking performance of the trained model was first assessed on the simulated tomograms containing proteins from the training set where the F1 picking score was calculated from the true positive, false positive, true negative, and false negative picks as described in Methods.

The median F1 picking score across all validation tomograms was 0.88 with a range from 0.76 to 0.98 (Supplementary Fig. 4a). Across all proteins in the training set, the median validation F1 picking score is 0.92 (Supplementary Fig. 4b). In rare cases, outlier scores were observed where specific proteins were unable to be picked across a range of sizes (Supplementary Fig. 4c). Closer inspection of these outliers revealed that in the simulated tomograms, each of these proteins display a particularly weak signal when compared to proteins of similar size (Supplementary Fig. 4d). In these cases, it appears that these proteins display a shape that is not recovered well during tomogram reconstruction by weighted back-projection. Despite this, picking on the validation tomograms demonstrated high accuracy for proteins across a wide array of shapes and sizes ranging from 30 kDa to 2.7 mDa.

### TomoTwin generalizes to unseen proteins

In order the assess the generalization of the general picking model to particles that were not in the training data set, we measured the picking performance with other, previously unseen proteins in a simulated tomogram with the reference-based workflow. We measured the F1 score of proposed particle locations against ground-truth boxes for seven proteins not included in the training data for which TomoTwin was therefore naïve (Fig. 2a,b). This assessment revealed that when trained on a set of 120 dissimilar proteins (Supplementary Fig. 3), the resulting model was able to locate all seven proteins accurately with a median F1 score of 0.82 despite a lack of previous training on these proteins (Fig. 2d). To measure the effect of training set size on generalization accuracy we performed this analysis on picking models trained on 20, 50, 100, and 120 proteins where we observed a logarithmic increase in generalization accuracy with the number of proteins in the training set (Fig. 2c). This high accuracy in locating novel proteins indicates a high generalization capability of TomoTwin.

**Fig. 2:**
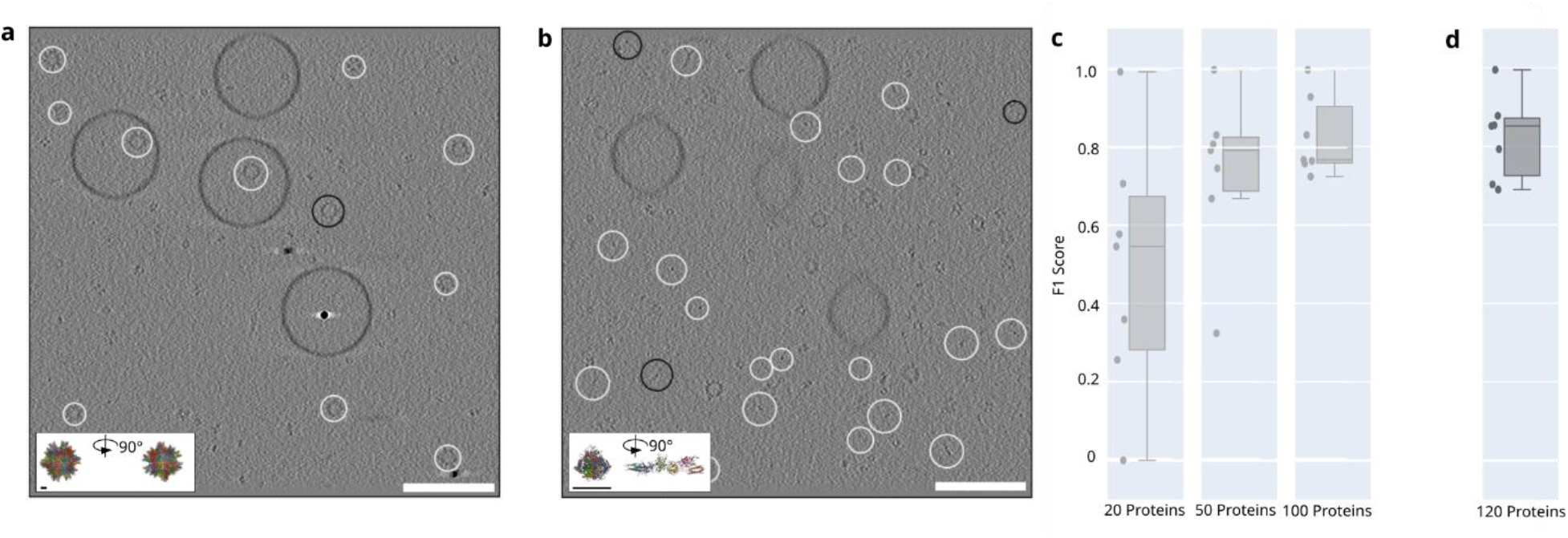
TomoTwin generalizes to novel proteins and locates them accurately. **a,** True positive selected particles (white) and false negative (black) of the largest protein 2DF7 (896 kDa) and **b,** the smallest protein 1FZG (142 kDa) in the generalization tomogram. The F1 scores are 0.99 and 0.88 for 2DF7 and 1FZG respectively. **c,** With increasing number of proteins used during training the mean F1 score on the generalization tomogram increased as well. The mean F1 scores are 0.49, 0.73, 0.82 for a model trained on 20, 50 and 100 proteins respectively. **d,** The model trained on the full training set of 120 proteins reached a mean F1 score of 0.82 but has the highest median F1 score of 0.85. White scale bar 100 nm, black scale bar 5 nm

As TomoTwin is trained entirely on simulated data, it is paramount to investigate its ability to pick proteins of interest in experimental tomograms. To evaluate this, cryo-ET was performed on a sample containing a mixture of three proteins, namely apoferritin^34^, the Type VI secretion effector RhsA from Pseudomonas protegens^34^, and the Tc toxin A component TcdA1 from Photorhabdus luminescens^35^ as well as liposomes (DOPC/POPC) (Fig. 3a). This mixture was chosen to create an environment with several proteins of different sizes as well as liposomes to mimic non-protein structures that may confound picking accuracy. Ten reconstructed tomograms were picked for apoferritin, RhsA, and TcdA1 using the pretrained general model of TomoTwin. In each case, the reference-based workflow was employed in which a target embedding was created by picking a single example of each protein as they are readily observable in tomograms with sufficient contrast. The target embedding from one tomogram was then applied across all tomograms in each dataset. Direct visualization of the picking similarity maps and final picking reveals high fidelity localization of each protein within the tomograms despite none of these proteins being included in the training set (Fig. 3b).

**Fig. 3:**
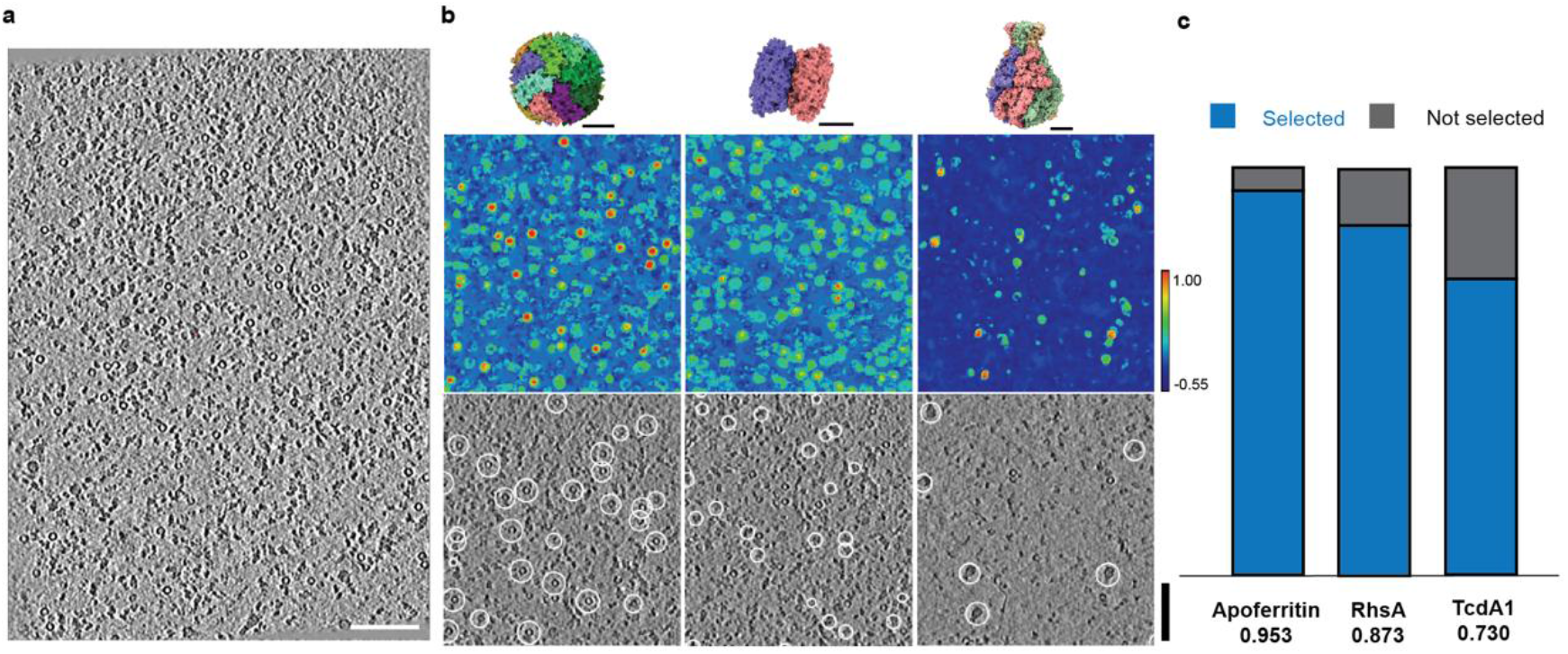
TomoTwin accurately localizes multiple proteins simultaneously in crowded tomograms. **a,** Representative slice of a tomogram containing a mixture of apoferritin, RhsA, and TcdA1; scale bar: 100 nm. **b,** Protein structure, cosine similarity map between tomogram and each target, and representative picking for apoferritin (PDB ID: 1DAT), RhsA (PDB ID: 7Q97), and TcdA1 (PDB ID: 6L7E) respectively. Scale bar for protein structures: 5 nm, scale bar for tomograms: 100 nm, color bar: -0.55 -1.00 **c,** Proportion of picked subvolumes contained within positive 2D classes. Total subvolumes picked: apoferritin: 848, RhsA: 577, TcdA1: 122.

As ground-truth particle coordinates are not available for experimental data, the accuracy of the picking was assessed by extracting subvolumes at the picked coordinates of each protein, projection of the 3D subvolumes to 2D using SPHIRE^36^, and performing 2D classification^37^ to evaluate the picked particles in a reference-free manner (Supplementary Fig. 5). For each protein of interest, the number of particles in the 2D classes displaying a high similarity to 2D classes of the protein previously determined by single particle analysis were recorded and represented as a percentage of the total number of particles picked (Fig. 3c). The high proportion of particles in all positive classes indicates that the picking is of high accuracy, confirming the visual impression of the picking result.

One of the principal advantages of cryo-ET is the ability to directly visualize proteins in their native cellular environments. Due to crowding of the cellular space and the poor contrast caused by thick specimens however, particle localization within a cellular environment presents a significant challenge. To assess its ability to locate particles in cellular tomograms, we applied TomoTwin to a dataset of tomograms containing Mycoplasma pneumoniae^38^ (EMPIAR 10499) (Fig. 4a). Using the TomoTwin general model, we picked 70S ribosomes in 65 tomograms with the reference-based workflow in which a reference was identified on one tomogram and used to generate a target embedding that was then applied to the entire dataset (Fig. 4b,c). To visualize the results, we extracted pseudo-subtomograms^39^ and performed 3D classification using a 70S ribosome cryo-EM structure (EMD 11650) lowpass filtered to 30 Å as a reference. As all 3D classes resemble ribosomes refined to ∼15 Å, it clearly indicates that TomoTwin also picks highly accurately in cellular tomograms (Fig. 4d).

**Fig. 4:**
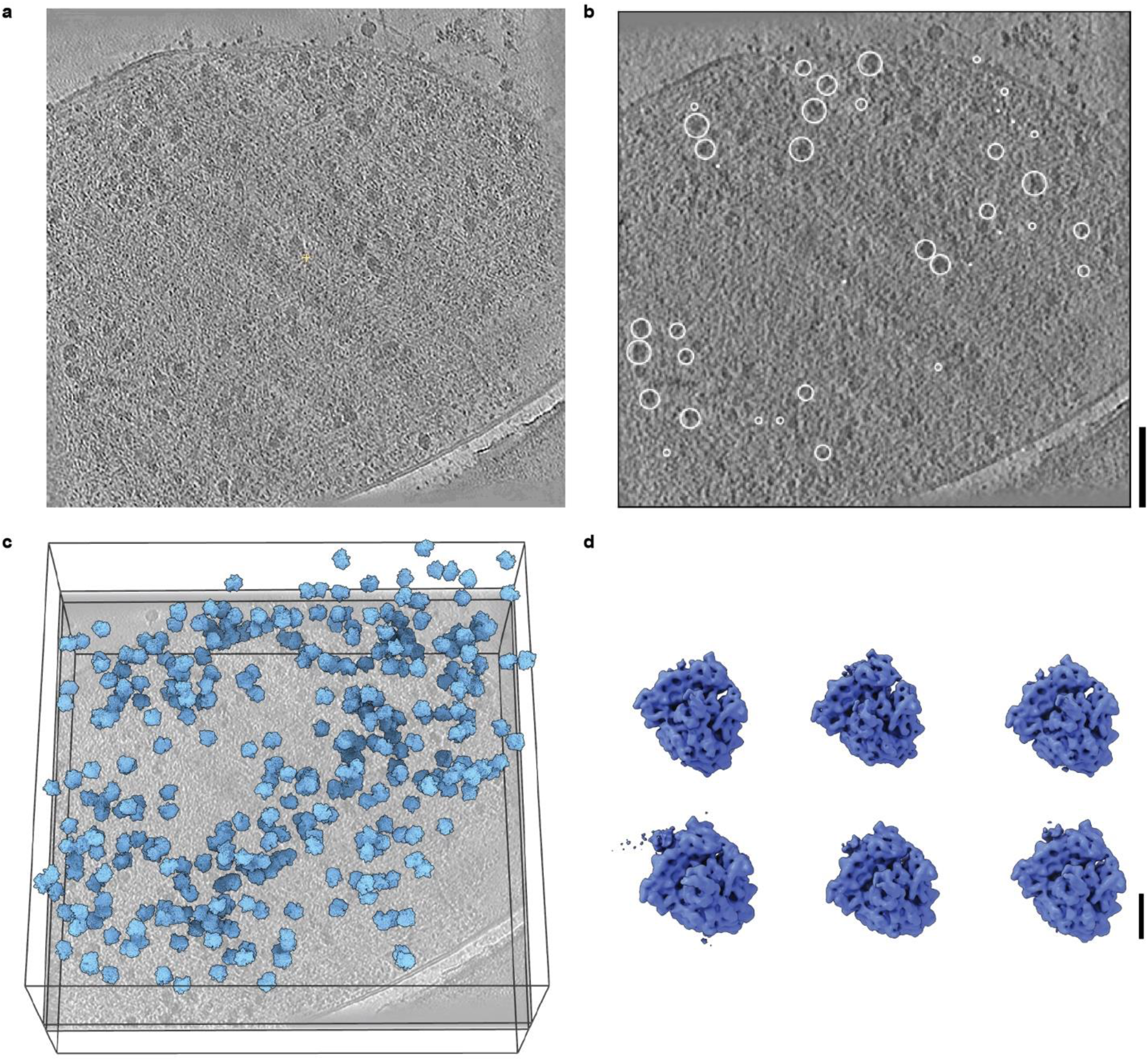
TomoTwin locates proteins in a cellular environment. **a,** Representative slice view of a tomogram containing Mycoplasma pneumoniae. b. Slice view highlighting positions of picked 70S ribosomes localized in 3D with TomoTwin. Scale bar 100 nm **c,** 3D representation of ribosome positioning within the tomogram, a represented slice is superimposed with 3D classes of ribosomes arranged according to their corresponding coordinates and orientation. **d,** 3D classes from 18,246 particles. Scale bar 10 nm.

### Structural Data Mining on the Embedding Manifold

The embedding feature of TomoTwin constructs a representation of a tomogram as a series of high-dimensional embeddings. These high-dimensional embeddings can be directly visualized by approximation on a 2D manifold (Fig. 5a,c). As a result of our deep metric learning-based approach, these representations contain a wealth of information about the contents of a tomogram where the distance between two subvolume embeddings directly correlates to the similarity of the 3-dimensional macromolecular shapes contained within. Typically, these representations contain a large mass corresponding to background noise, or a particularly prominent feature of the tomogram volume as well as additional well-defined clusters corresponding to different shapes such as proteins, membranes, or fiducials. By directly visualizing these representations on a 2D manifold, the clustering workflow of TomoTwin allows interactive, structural data mining of tomograms, where clusters of subvolumes on the embedding manifold are used to locate different macromolecular populations within the tomogram.

**Fig. 5:**
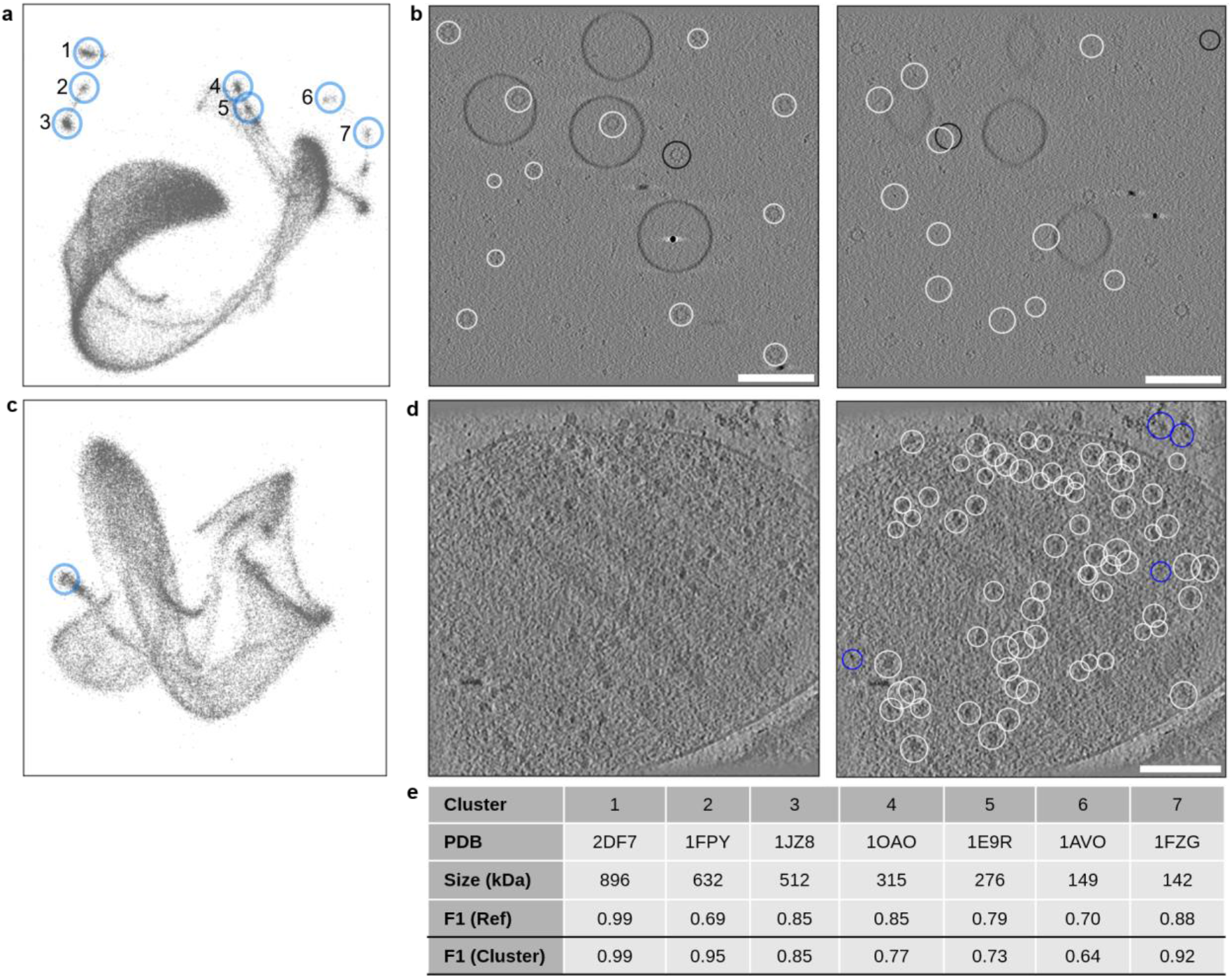
TomoTwin enables structural data mining on the embedding manifold. **a,** Highlighted clusters of all 7 proteins on the generalization tomogram 2D manifold approximation. **b,** Respective particle locations from cluster 3 and 5 which corresponds to the proteins with PDB ID 2DF7 (left) and 1FZG (right). White are true-positive picks and black false-negative. In both cases there were no false-positive selections. **c**, 2D manifold approximation of the embedding space of a tomogram containing Mycoplasma pneumoniae (EMPIAR 10499). Highlighted is the manual selected cluster which corresponds to the 70S ribosome. **d,** Using the cluster center for picking identified all ribosomes previously selected by the reference-based picking (white) with a few reference-only selections (blue). **e,** F1 scores for the individual clusters in comparison with the F1 scores for reference-based picking. On average the clustering performed slightly better (0.84 vs 0.82 mean F1 score). However, for some individual proteins the difference was larger (e.g. cluster 7). Scale bar 100 nm

To evaluate the accuracy of clustering-based picking quantitatively, we again utilized our simulated generalization tomogram where we evaluated the results of the clustering-based picking of each protein against the ground-truth coordinates with the F1 picking score as well as directly comparing it against the reference-based workflow (Fig. 5b,e). The clustering-based picking identified each protein with high accuracy across a range of sizes. Notably, it outperforms the reference-based workflow for glutamine synthetase^40^ (PDB ID: 1FPY) indicating that this workflow provides complementary advantages to the reference-based workflow. Additionally notable in the manifold projection of the representation map is the fact that individual protein clusters are globally organized by size, with the three largest protein clusters located in one area of the map, clusters for medium sized proteins in another, and the clusters for the two smallest proteins located furthest away from those of the large proteins, demonstrating that the model accurately represents complex similarity relationships in terms of protein structures as distance in the embedding space (Fig. 5a).

We additionally compared the clustering-based picking workflow directly against the reference-based approach for the cellular tomograms containing M. pneumoniae (Fig. 5c). Examining the representation maps of these tomograms, several clusters are visible. One of which, when picked, produces accurate particle locations for 70S ribosomes nearly identical to those produced by the reference-based approach once again underlining the robustness of both workflows (Fig. 5d).

## Conclusion

Despite offering the potential to study proteins in their native, cellular environment in unprecedented detail, it remains that, presently, only a select few proteins have been successfully studied in detail by cryo-ET with STA. In part, this is because with increased cellular context, the formation of macromolecular complexes, and poorer contrast caused by thicker specimens, comes the challenge of picking individual proteins for subsequent subtomogram averaging. To assist in this crucial particle picking step, we developed TomoTwin, a robust, first in class general picking model for cryo-electron tomograms based on deep metric learning. TomoTwin allows users to identify proteins in tomograms *de novo* without manually creating training data or retraining the network each time a new protein is to be located.

The innovation landscape for algorithm development in both cryo-EM and cryo-ET bears a heavy emphasis on automated processing for increased data throughput^26, 37, 41– 43^ . With its highly generalizable picking model, TomoTwin is the first tool based on deep learning that can be readily integrated with high throughput tomogram reconstruction and STA workflows. Additionally, when combined with unsupervised cluster detection algorithms^44^, the clustering workflow of TomoTwin paves the way for unsupervised STA analysis on a whole-tomogram level (Supplementary Fig. 6).

TomoTwin is a robust, open-source tool for particle localization in cryo-electron tomograms. The code used to develop and train TomoTwin as well as the general picking model and tools to use it for generalized particle picking are available at https://github.com/MPI-Dortmund/tomotwin-cryoet with future updates including extensive user documentation available soon.

## Methods

### Training Data Generation

TomoTwin was trained on 123 data classes comprised of subvolumes of 120 different proteins, membranes, noise, and fiducials from simulated tomograms. To ensure that TomoTwin is trained on the most diverse set of proteins possible, 108 proteins were selected from the PDB with sizes ranging from 30 kDa to 2.7 mDa and the cross correlation between pairs of 10 Å low-pass filtered maps of each protein was calculated (Supplementary Figure 3). Any protein with a high similarity (greater than 0.6) to another protein in the training set was marked for replacement. Additionally included were the data from the 2021 SHREC competition including 12 proteins^18^ to yield a total of 120 proteins for training. A training/validation split was achieved with 800 subvolumes for each data class in the training set and 200 in the validation set, yielding a total training set size of 98,400 subvolumes and a validation set size of 24,600 subvolumes.

### Tomogram simulation

Tomogram simulation was done using TEM Simulator^45^ which calculates the scattering potential of individual proteins and places them in definable positions within the volume. The output of the simulation is a tilt series which is then reconstructed using IMOD^46^. A configuration file was generated with properties for the electron beam, optics of the microscope, the detector, the tilt geometry and the sample volume. The default detector was adjusted to reflect the MTF curve of a modern Gatan K3 Camera with a quantum efficiency of 0.9. The detector size was set to 1024×1024 with a pixel size of 5 micrometer. The magnification was set to 9800, the spherical aberration and chromatic aberration were adjusted to 2.7 mm and 2 mm respectively to mimic popular modern TEMs. A condenser aperture size of 80 micrometer was chosen. For each tomogram the defocus value was randomly chosen between -2.5 µm and -5 µm. A tilting scheme of -60° to +60° with a step size of 2° was used. To simplify and streamline the simulation we wrote a set of open-source programs called “tem-simulator-scripts” (https://github.com/MPI-Dortmund/tem-simulator-scripts). They contain scripts that require as input the PDB files to be simulated and the number of particles to simulate per PDB. The program then generates reconstructed tomograms as they were used for this study using the following pipeline:

1. Generation of densely packed random particle positions within the volume where individual particles do not overlap.
2. Generation of an occupancy map - a volume where each voxel is labeled according the protein identity.
3. Generation of fiducial maps.
4. Generation of vesicle maps.
5. Generation of the configuration file for TEM-simulator
6. Simulation of the tiltseries using TEM-simulator.
7. Alignment and reconstruction using IMOD.

However, all steps can also be carried out individually to have full control over all parameters.

Using this procedure, we simulated 11 sets of proteins. The sets contain in total 108 different proteins with each set covering proteins of various sizes. For each set we simulated 8 tomograms of size 512×512×200 voxels with a pixel size of 1.02 nm and varying protein density. For tomogram 1, 2 and 8, 150 particles per protein were generated, for tomogram 3 and 4, 125 particles per protein, for tomogram 5 and 6, 100 particles per protein and for tomogram 7, 75 particles per protein. Tomograms 1-7 were used for training and tomogram 8 for validation. The generated tomograms used in this study with all meta-data are publicly available^47^. These simulated data were used to construct the training and validation sets^48^ to evaluate network training, particle localization, and model generalizability.

### Convolutional Network Architecture

To encode volumetric cryo-ET data as embedding vectors in a high-dimensional space, TomoTwin employs a 3D CNN consisting of five convolutional blocks followed by a head network (Supplementary Fig. 1a). Each convolution block consists of two 3D convolutional layers with a kernel size of 3×3×3. Each convolutional layer is followed by a normalization layer and a leaky rectified linear (ReLU) activation function. In the first convolutional layer of each convolutional block, the number of output channels is twice the input channels and in the second convolutional layer the number of output channels matches the output from the previous layer. Max pooling is performed with a kernel size of 2×2×2 after the first convolutional block and adaptive max pooling to a size of 2×2×2 is performed after the final convolutional block. As a result, when provided with a 37×37×37 subvolume with 1 channel as a normalized, 37×37×37×1 array, the convolutional blocks transform the input to a 2×2×2×1024 feature vector which is then fed to the head network. In the head network, the feature vector is first flattened channel-wise before being subject to a dropout layer and then passed through a series of fully-connected layers that transform the flattened vector to a 1-dimensional, 32-length feature vector. Finally, this feature vector is L2-normalized to yield an output embedding vector for the subvolume.

### Triplet Generation

TomoTwin is trained on triplets of subvolumes consisting of an anchor volume A, a positive volume P, and a negative volume N (Supplementary Fig. 1b). Each subvolume is assigned to a data class corresponding to the macromolecule contained within and has a size of 37×37×37 voxels. Triplets are constructed where A and P are sampled from the same data class and N from a different data class. Given a distance function D and an embedding function f, the triplet loss is defined as:

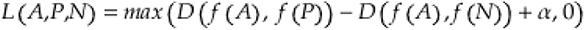

where the hyperparameter α is the margin value. As distance function D we use cosine similarity which is defined as

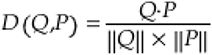

where Q and P are arbitrary embedding vectors, • is the dot product and ‖·‖ the length of the vector. During training, triplets are generated by online semihard triplet mining wherein a batch of subvolumes are embedded and triplets generated automatically with the negative subvolume embedding being selected from those only with a distance to the anchor greater than the positive subvolume embedding but not greater than a margin *α_miner_*:

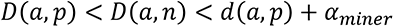

Where a, p and n are the embedding vectors of the anchor, positive and negative respectively and *α_miner_* is the margin of the miner.

### Training of the General Picking Model

Training of the 3D CNN was performed for 600 epochs using an adaptive moment estimation (ADAM) optimizer^49^. The model from the epoch with the best F1 score on the subvolumes in the validation set was further evaluated in the localization and generalization tasks and used as the general picking model.

### Data augmentation

To prevent overfitting during training and to improve generalization of the model, online data augmentations were applied to each normalized volume before its embedding was calculated including rotation, dropout, translation, and the addition of noise. For the rotation augmentation, subvolumes were rotated by a random angle in the X-Y plane but not X-Z or Y-Z to prevent reorientation of the missing wedge. In the dropout augmentation, a random portion between 5 and 20% of the voxels were set to the subvolume mean value. In the translation augmentation, the subvolume was shifted by 1-2 pixels in each direction. The addition of noise augmentation added Gaussian noise with a randomly chosen standard deviation between 0 and 0.3 to the subvolume.

### Hyperparameter optimization

The training of modern convolutional neural networks involves the selection of many hyperparameters, some of these choices affect the architecture while others affect the learning process itself. While some heuristics exist to guide hyperparameter selection, finding a combination of settings that maximize the utility of a machine learning tool by hand quickly becomes intractable. Optuna^50^ was applied to explore the hyperparameter search space and identify an optimized set of parameters for training . Models were trained on a subset of the training data for 200 epochs and the F1 score calculated on the validation set after each epoch. Pruning was performed after 50 epochs for training runs with an F1 score lower than the global median. In total, searches were applied for the hyperparameters of learning rate, dropout rate, optimizer, batch size, weight decay, size of the first convolution kernel, number of output layer nodes, online triplet mining strategy (semihard^51^, easyhard^52^ , none), normalization type (group norm^53^, batch norm^54^), loss function (TripletLoss^31^, SphereFace^55^, ArcFace^56^), and loss margin (Supplementary Fig. 7).

Most notably and unexpectedly, the type of normalization applied during training was the largest overall affecter of performance with group normalization^53^ outperforming the more common batch normalization^54^ strategy (Supplementary Fig. 7b). Additionally noted was the increased performance of a standard triplet loss function over the theoretically superior SphereFace^55^ and ArcFace^56^ loss functions (Supplementary Fig. 7c). These findings underpin the necessity to explore a wide range of hyperparameters during training as heuristics alone are not enough to guide optimal hyperparameter selection for the training of modern convolutional neural networks.

### Particle picking workflow with the general model

For each dataset picked with the general model, first all tomograms were embedded. To achieve this, the tomograms were subdivided into a series of overlapping 37×37×37 subvolumes with a stride of 2 voxels. For the reference-based workflow, a random particle for each protein of interest was selected as reference and embedded to generate a target embedding. The tomogram and target embeddings were provided to TomoTwin Map which calculated the distance matrix between each target embedding and each subvolume embedding from the tomogram and returned this along with a similarity map for each target embedding. This matrix was then provided to TomoTwin Locate which identified areas of high confidence as target locations using a region-growing based maximum detection procedure followed by non-maxima suppression. The returned candidate positions were then subject to confidence and size thresholding with TomoTwin pick to produce final coordinates for each protein of interest.

### Evaluation of simulated data

The performance of particle localization was calculated from three metrics: recall, precision, and, the harmonic mean of the two, the F1 score which are defines as:

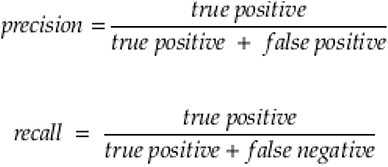

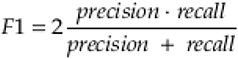

Selected particle locations counted as true positives if the intersection over union (IOU) of the box of the selected particle location and the ground truth box was greater than 0.6. The IOU is defined as the ratio of the intersecting volume of two bounding boxes and the volume of their union.

The particle localization accuracy of the trained model was assessed for each tomogram in the validation set (Supplementary Fig. 4a). To test model generalization, the localization task was performed on a tomogram containing 7 proteins not included in the training set for which TomoTwin was therefore naïve (Fig. 2).

### Clustering

For clustering analysis, a random sample of 400,000 embeddings from the high-dimensional tomogram embeddings were fit to a uniform 2D manifold with Uniform Manifold Approximation (UMAP) with GPU-acceleration provided by the RAPIDS package^57^. The UMAP model was used as the basis to transform the entire tomogram embeddings and the results plotted (Figure 5a,c). Clusters were identified by eye and selected by drawing a closed shape containing the desired points. The enclosed points were then traced back to their original high-dimensional embeddings and the average embedding of them was calculated. This average embedding was then used as a target embedding for classification, localization, and picking in the same manner as for the reference-based workflow.

### Preparation of Experimental Samples

The components of the mixture were either thawed from long-term storage at -80 °C or freshly prepared. *Photorhabdus luminescens* holotoxin was expressed, purified and the holotoxin formed as described previously^58^ and used at a stock concentration of 0.49 mg/mL. RhsA from Pseudomonas protegens was expressed and purified as described previously^34^ and used at 4 mg/mL concentration. Liposomes were prepared by extrusion. 4 mg/mL of each POPC (1-palmitoyl-2-oleoyl-glycero-3-phosphocholine, Avanti Polar Lipids) and DOPS (1,2-dioleoyl-sn-glycero-3-phospho-L-serine, Avanti Polar Lipids) were mixed in buffer (50 mM Tris, pH 8, 150 NaCl, 0.05% Tween20) and after brief sonication (1 min in water bath) and three cycles of freeze-thawing (−196 °C and 50 °C), the liposome solution was passed 11 times through a polycarbonate membrane with a 400 nm pore size in a mini extruder (Avanti Polar Lipids). Total lipid concentration was diluted with buffer to 0.16 mg/mL. The freeze-dried content of one vial Tobacco mosaic virus (TMV) (DSMZ GmbH Braunschweig, Germany, PC-0107) was solved in 1 mL buffer and diluted 500 times as working solution. The Apoferritin (ApoF) plasmid was a kind gift by Dr. Christos Savva (Midlands Regional Cryo-Electron Microscopy Facility). Expression and purification of ApoF was optimized based on the protocol described earlier^59^ and final concertation of frozen stock was 3 mg/mL.

Different ratios of the mixture were prepared and then examined after vitrification using cryo-EM. For cryo-ET, a ratio of 1:2:2:20:10 (TMV:Apoferritin:Liposomes:TcToxin:RhsA) was chosen.

### Grid Preparation

Grids were prepared using a Vitrobot Mark IV (Thermo Fisher Scientific) at 4 °C and 100% humidity. 4 µL of the freshly prepared mixture were applied to glow-discharged (Quorum GloQube) R1.2/1.4 Cu 200 (Quantifoil) grids. After blotting (3.5 s at blot force -1, no drain time) the specimen was vitrified in liquid ethane.

### Cryo-ET

Grids of different mixing ratios were screened using a Talos Arctica electron microscope (Thermo Fisher Scientific) equipped with a X-FEG and Falcon 3 camera. Small datasets of 100-200 images were collected using the software EPU (Thermo Fisher Scientific). The best specimen was transferred to a Titan Krios G3 electron microscope equipped with X-FEG. Images were recorded on a K3 camera (Gatan) operated in counting mode at a nominal magnification of 63,000, resulting in a pixel size of 1.484 Å/pix. A Bioquantum post-column energy (Gatan) was used for zero loss imaging with a slit width of 20 eV. Tilt series were acquired using SerialEM^60^ with the Plugin PACEtomo^61^ and with a dose symmetric tilt scheme^62^ from 60° to 60° with a step size of 3°. Each movie was collected as an exposure of 0.2 seconds subdivided into 10 frames. Frames were then exported to Warp 1.0.9^26^ for motion correction, CTF estimation and generation of tilt series. Tilt series were aligned with patch tracking and tomograms reconstructed by weighted back-projection in IMOD^47^ with a pixel size of 5.936. Tomograms were scaled by Fourier shrinking to 10 Å/pix for embedding with TomoTwin.

Raw frames of M. pneumoniae cells were downloaded from EMPIAR (EMPIAR-10499). Motion correction and CTF estimation were performed in Warp 1.0.9 which was then used to generate tilt series. These tilt series were aligned with patch tracking and tomograms reconstructed by weighted back-projection in IMOD with a pixel size of 6.802 Å/pix. Tomograms were then scaled by Fourier shrinking to 13.6 Å/pix for embedding with TomoTwin.

### Evaluation of experimental data

For tomograms from samples prepared in-house, coordinates of particles identified with TomoTwin were scaled to a pixel size of 5.936 to match the originally reconstructed tomograms. The tomograms were imported and these coordinates were used to extract subtomograms in Relion 3.0^37^. For reference-free analysis, 3D subtomograms were projected to 2D with SPHIRE^36^ and then used for 2D classification. For tomograms attained from EMPIAR, coordinates of particles identified with TomoTwin were scaled to a pixel size of 6.802 Å/pix to match the originally reconstructed tomograms. The tomograms were imported and coordinates were imported and used to reconstruct pseudo-subtomograms in Relion 4.0^40^. A reference was created from a 70S ribosome (EMD-11650) by lowpass filtering to 30 Å and then scaling the pixel size to 6.802 Å/pix. This reference was used for 3D classification with the pseudo-subtomograms in Relion 4.0.

### Hardware

Two computational setups were utilized for calculations, a distributed computing system and a local workstation. The distributed computing system consisted of the Max Planck Gesellschaft Supercomputer ‘Raven’ using up to 30 Nvidia A100 GPUs, where each GPU has 40 GB memory. Each process had 18 cores of Intel Xeon IceLake-SP 8360Y processors and 128GB system memory available. The local workstation consisted of a local unit equipped with a Nvidia Titan V (12 GB memory) GPU and a Intel i9-7920X CPU with 64 GB system memory.

Hyperparameter optimization was done in parallel for 7 days one the distributed computing setup and embeddings were calculated on this set up as well using 2 GPUs. In all cases a box size of 37 and stride of 2 were used for embedding.

The inhouse workstation was used for miscellaneous tasks and for calculating timings using 2 GPUs.

### Timings

The calculation of the embeddings is the only function of TomoTwin requiring significant processing time. To measure this, we embedded our largest experimental tomogram (608×855×148 after Fourier shrinking) on a local workstation and a distributed computing system. Using 2 GPUs, tomogram embedding took 80 minutes for the local setup and 30 minutes for the distributed setup, corresponding to the total time to pick all proteins of interest per tomogram on each setup.

## Data Availability

All simulated tomograms used in this study are available here: https://doi.org/10.5281/zenodo.6637357.

The extracted subvolumes used to train and evaluate the performance of TomoTwin are available at: https://doi.org/10.5281/zenodo.6637456.

The TEM-Simulator-Scripts package used for automated tilt-series simulation and reconstruction is available at: https://github.com/MPI-Dortmund/tem-simulator-scripts.

TomoTwin is available under an open-source license at: https://github.com/MPI-Dortmund/tomotwin-cryoet.

## Author contributions

Conceptualization: T.W. and S.R.;

Software implementation: T.W., G.R., M.S.;

Software – Testing: G.R., T.W., M.S.;

Formal analysis: G.R., T.W.;

Supervision: S.R.;

Writing – original draft: G.R., T.W.;

Writing – review and editing: G.R., T.W., and S.R.;

Funding acquisition: S.R.;

## Acknowledgements

We thank D. Prumbaum for collecting cryo-ET data collection, P. Günther for providing RhsA and liposomes, P. Njenga Ng’ Ang’ A for providing TcdA1, and K. Vogel-Bachmayr for purifying apoferritin, C. Savva for providing the apoferritin plasmid and A. Prajica for support in figure and logo creation. The work was supported by the Max Planck Society (to S.R.).

---

**Supplementary Fig. 1:**
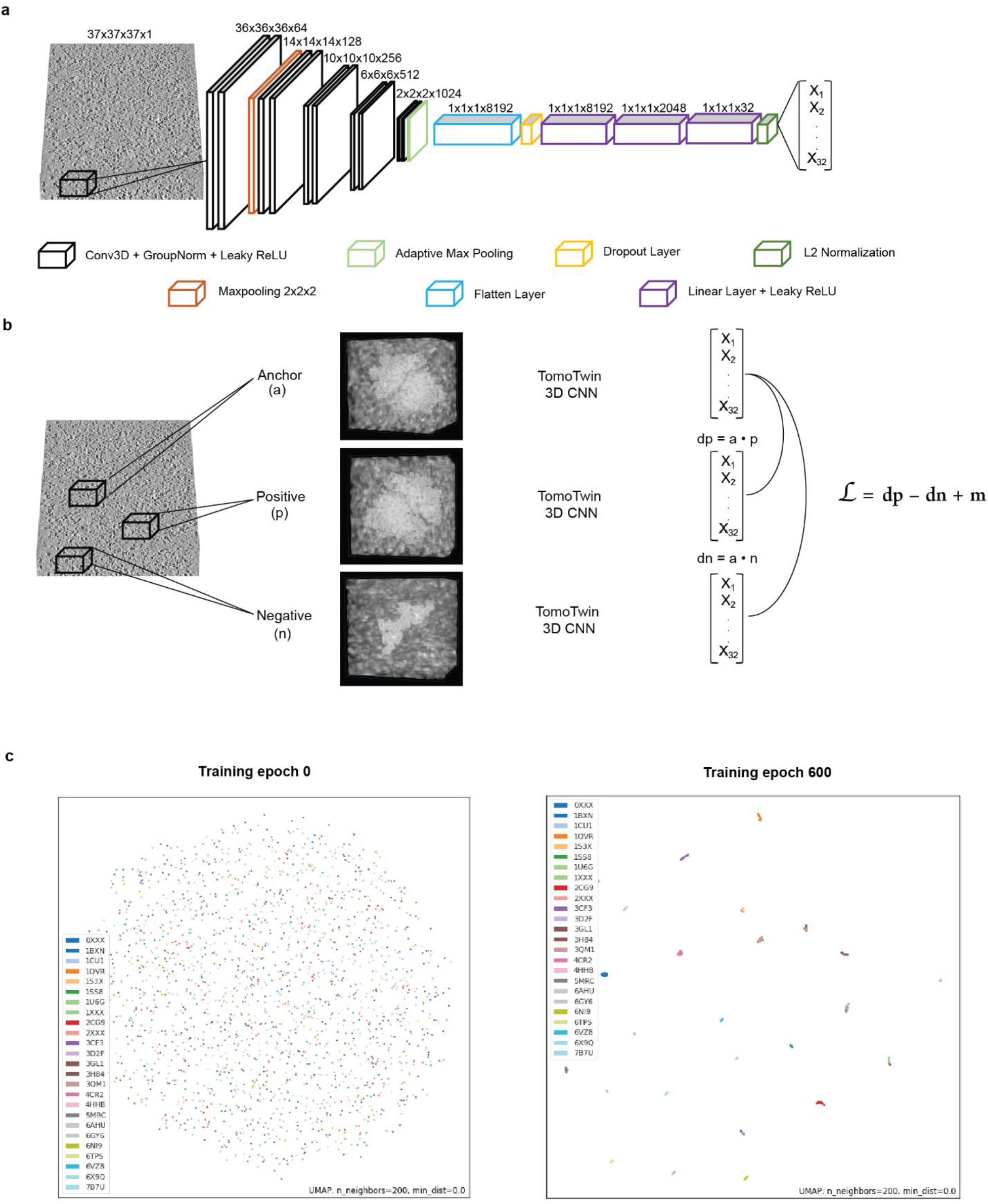
TomoTwin convolutional architecture and metric learning strategy. **a,** Architecture of 3D convolutional network utilized by TomoTwin to translate 3D real space tomogram subvolumes into embedding vectors for deep metric learning. **b,** Overview of the deep metric learning training scheme employed by TomoTwin wherein data triplets are constructed of anchor, positive, and negative subvolumes. The triplets of subvolumes are each convolved by the 3D CNN of TomoTwin and the resulting embedding vectors are used to calculate the distance metrics implicit in the triplet loss function. **c,** Uniform manifold approximation of protein subvolume embeddings colored according to protein PDB code from TomoTwin 3D CNN in first training epoch and best model after 600 training epochs.

**Supplementary Fig. 2:**
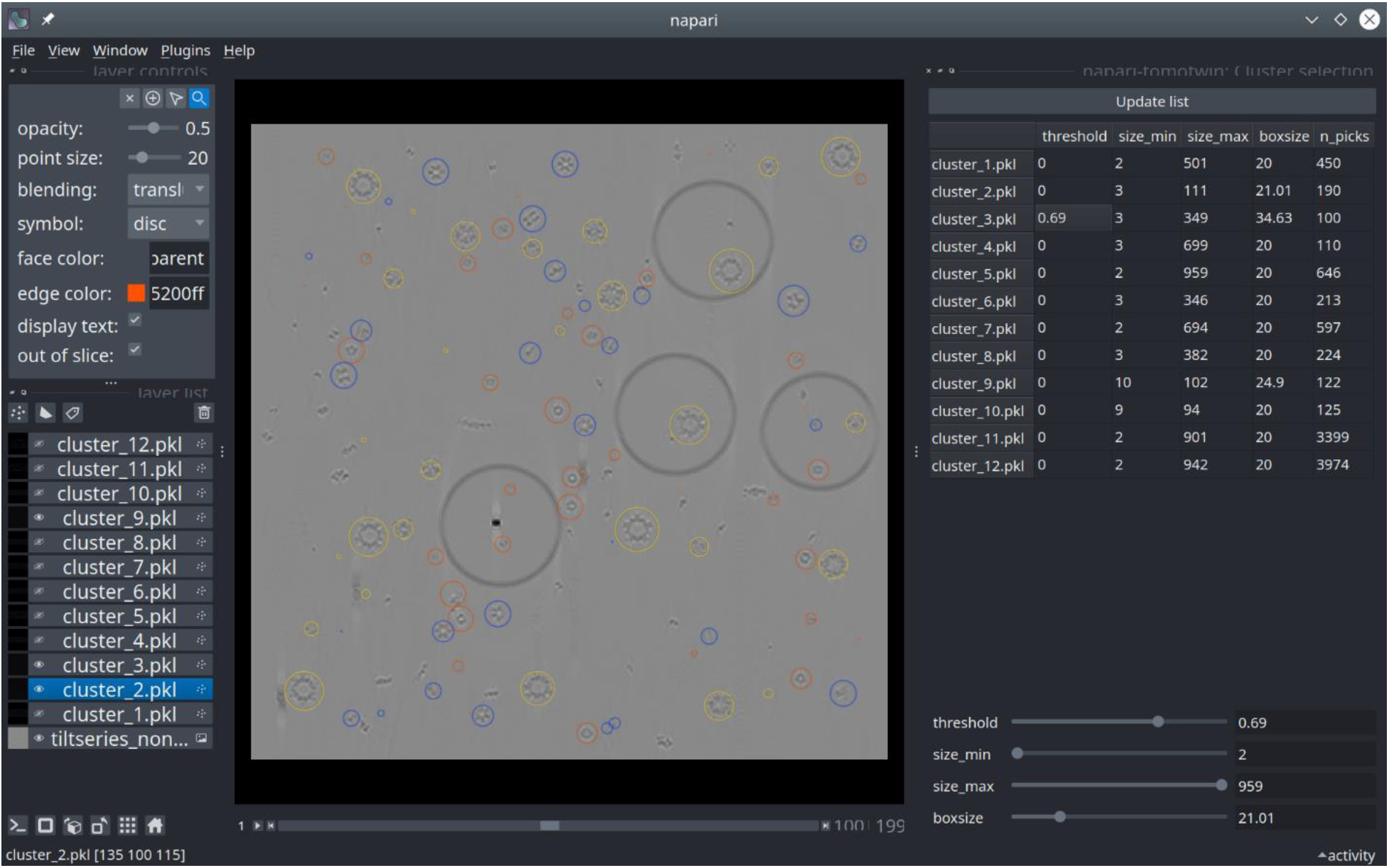
Graphical user interface of TomoTwin implemented as a Napari plugin. Visualization of protein picks in simulated generalization tomogram identified by the clustering workflow. Picks for 3 out of 12 clusters are shown as spheres. The lefthand panel allows users to adjust various visualization settings for the tomogram including 3D viewing as an isosurface. The righthand panel allows users to filter picks for each cluster according to similarity threshold, minimum and maximum size, and adjust the box size for viewing.

**Supplementary Fig. 3:**
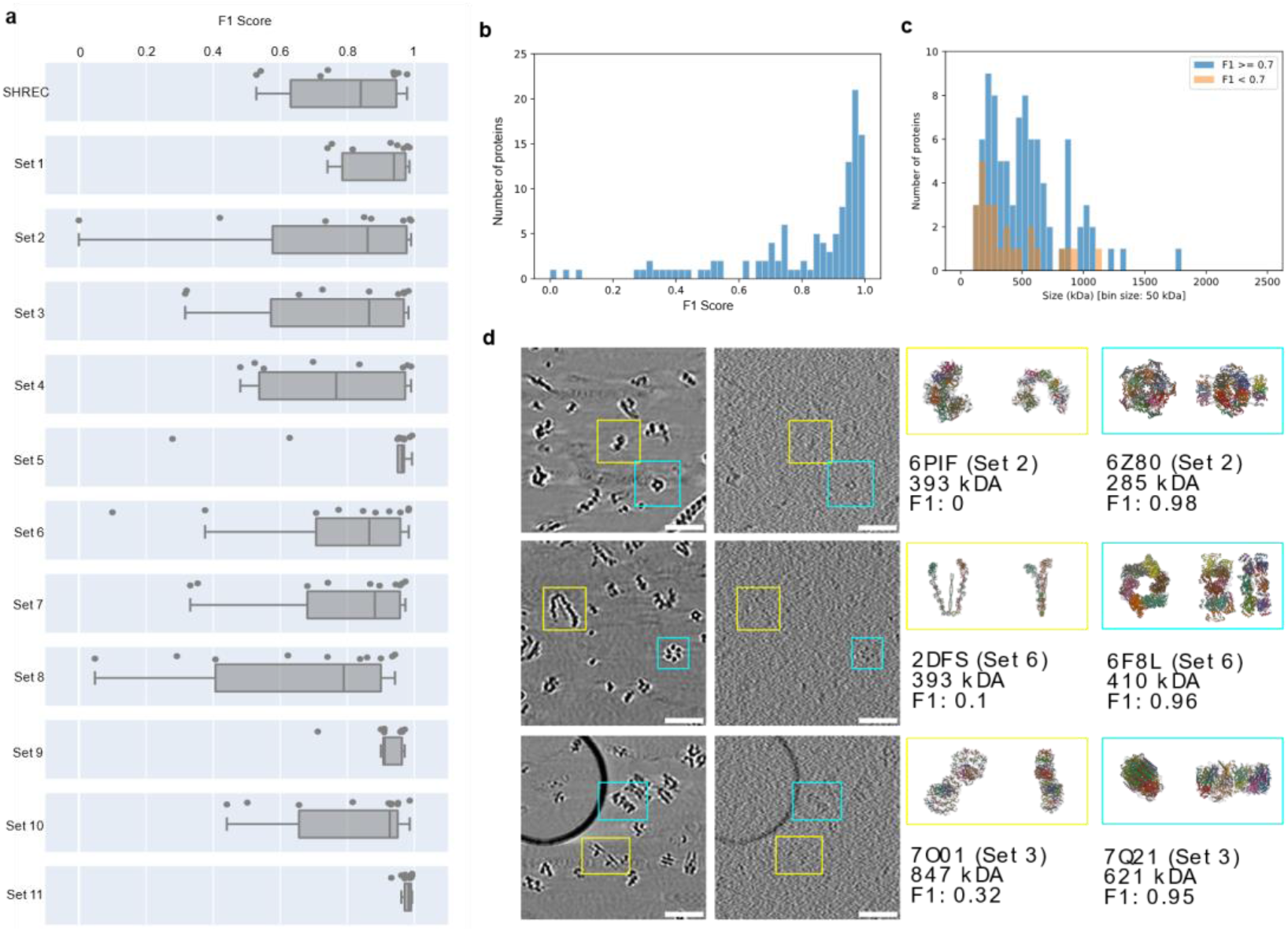
TomoTwin identifies proteins with high accuracy by using single particle subvolumes as reference. **a,** F1 scores of TomoTwin on the validation tomograms. The median F1 score of the individual sets is most often above 0.8 and not lower then 0.76. **b,** The overall distribution of F1 scores with a median of 0.92. However, a tail of proteins with low F1 scores can be seen. **c,** Size distribution of particles that show good F1 scores (F1>=0.7) and those with rather low F1 scores (F1 < 0.7). **d,** Examples of proteins of similar size with low (yellow) and high (cyan) F1 score. On the left side the individual particles are depicted in a noisy and noise free reconstruction, respectively. On the right side, the respective structures, F1 scores and sizes are shown. It can be seen that the proteins which were not identified properly by TomoTwin have a lower contrast than the other proteins. Scale bars 100 nm.

**Supplementary Fig. 4:**
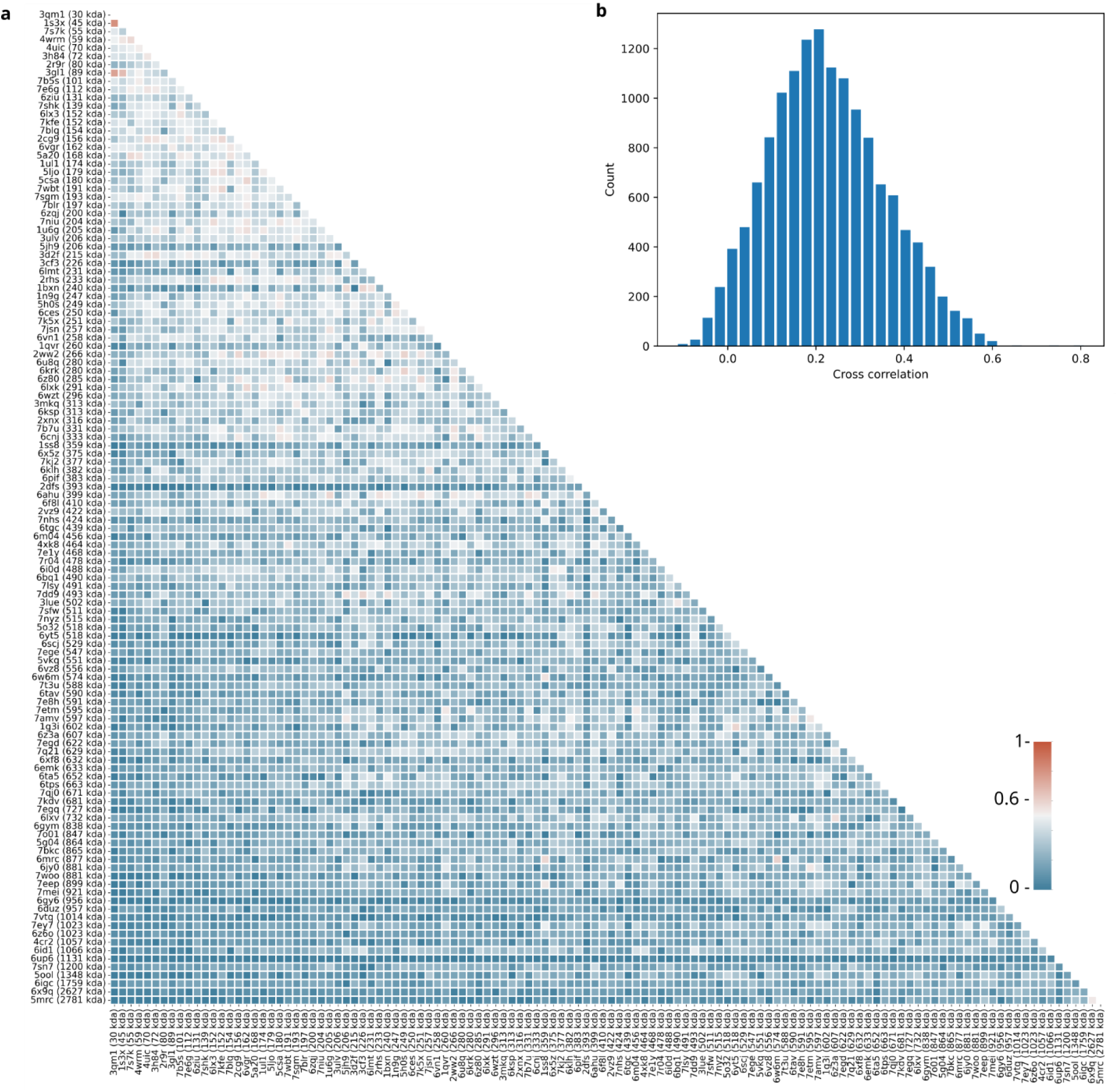
Characterization of the training data set. **a,** Pairwise cross-correlation matrix for all 120 proteins sorted by size. Cross-correlations were calculated by converting the individual PDBs to density maps with a pixel size of 1 nm, aligning them pairwise with EMAN2 and calculating the cross-correlation of the aligned pairs. To maximize the value for training, we selected proteins so that all pairs except 3 have a cross correlation value below 0.6. The three pairs with higher correlation are from the SHREC dataset and were not simulated by us. Higher correlation values are more likely for smaller proteins. **b,** Histogram of the pairwise cross correlation values. The mean cross correlation value is 0.22 with a standard deviation of 0.13.

**Supplementary Fig. 5:**
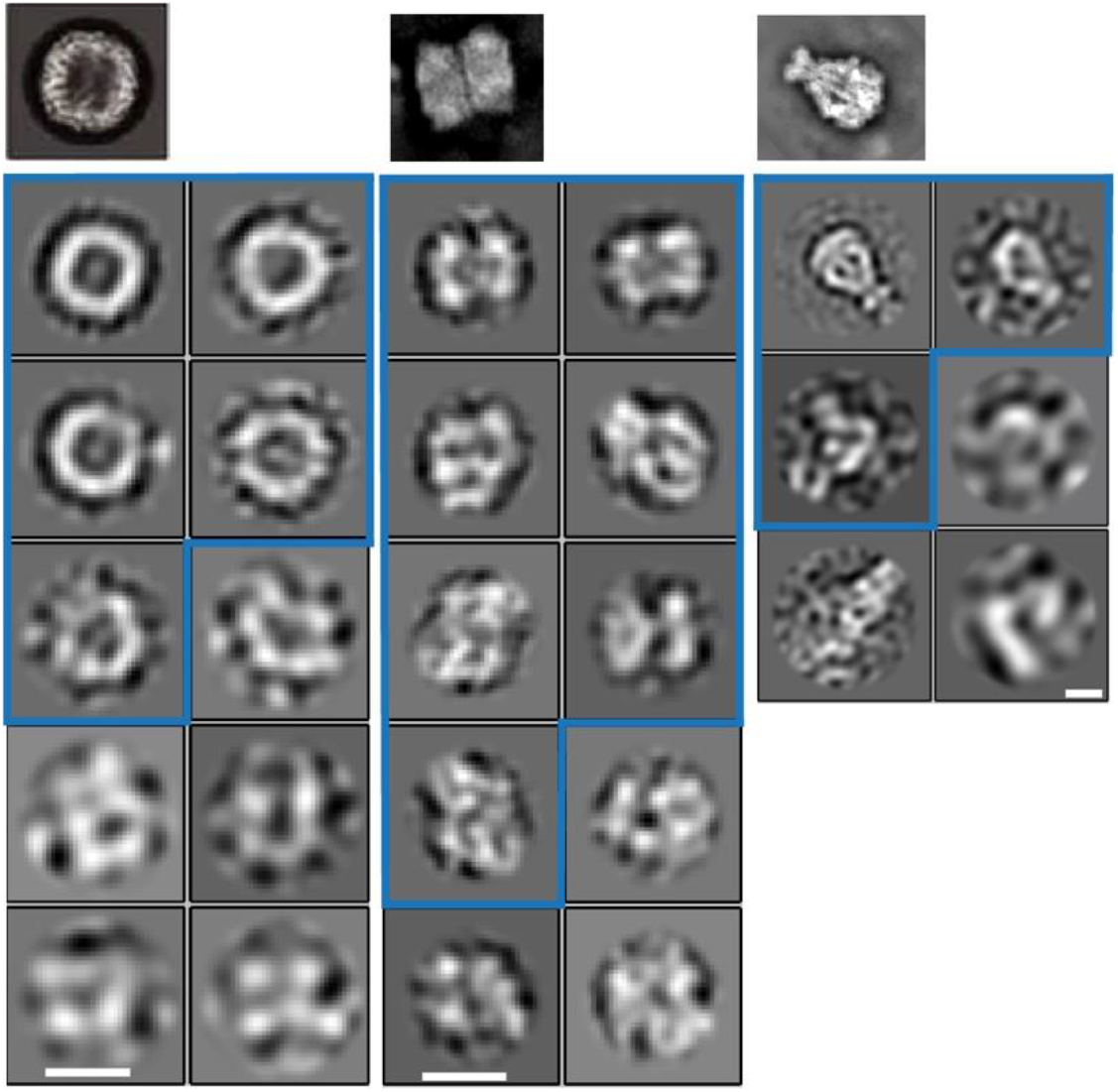
2D classes of proteins identified in a mixed tomogram. Example 2D classes from previous studies by single particle analysis of apoferritin^59^, RhsA^34^, and TcdA1^58^ respectively; 2D class averages of TomoTwin picked subvolumes after projection to 2D. Classes outlined in blue were judged to be positive classes by expert inspection, indicating that they contain particles of the appropriate protein. Scale bar: 5 nm

**Supplementary Fig. 6:**
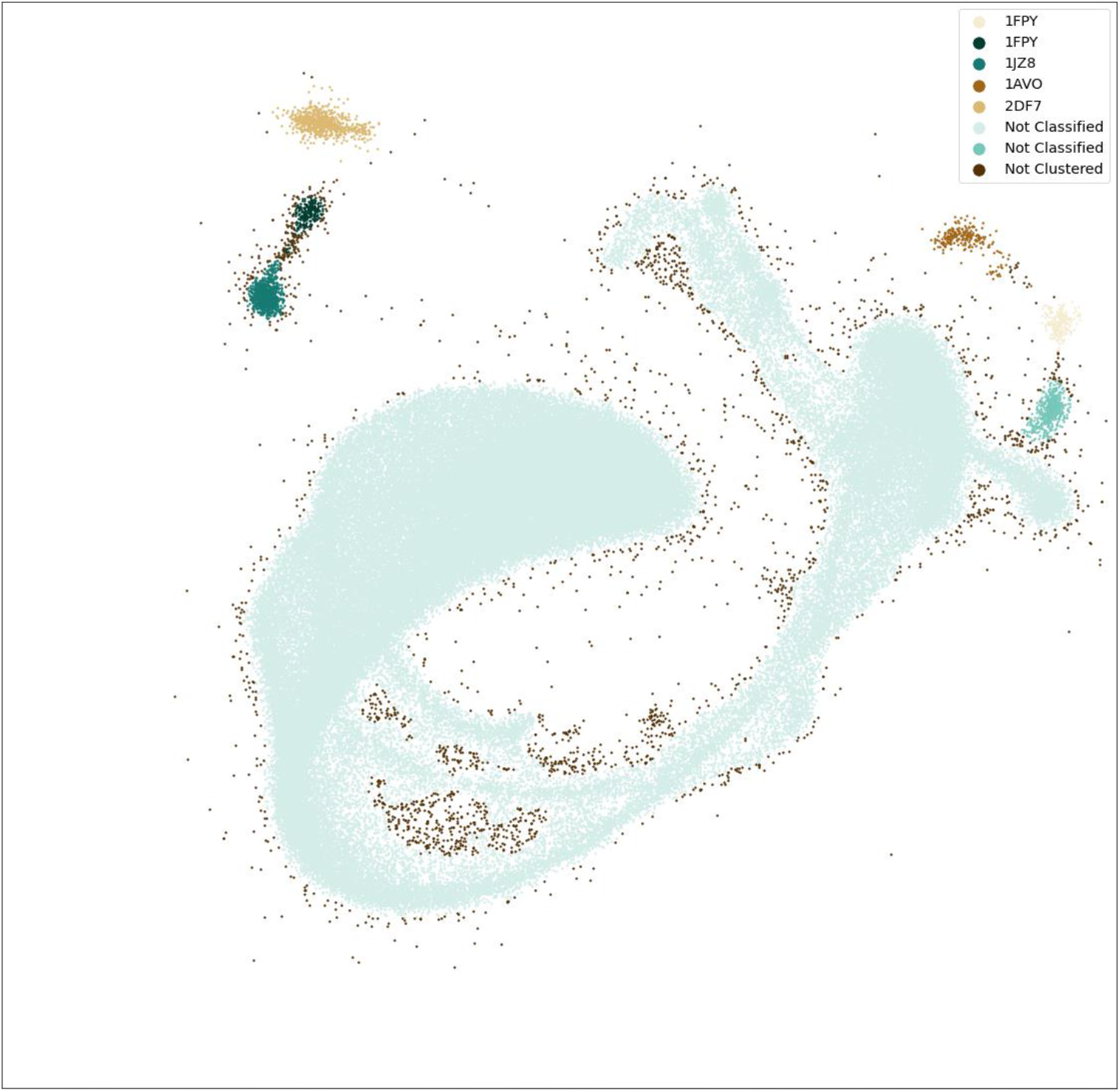
Automated identification of clusters of interest using HDBSCAN. A subset of the approximated manifold of Figure 5a was used to run density-based clustering which located 5 out of 7 clusters of interest in an unsupervised. R implementation of HDBSCAN was run with a min_samples of 50 and a minimum cluster size of 50.

**Supplementary Fig. 7:**
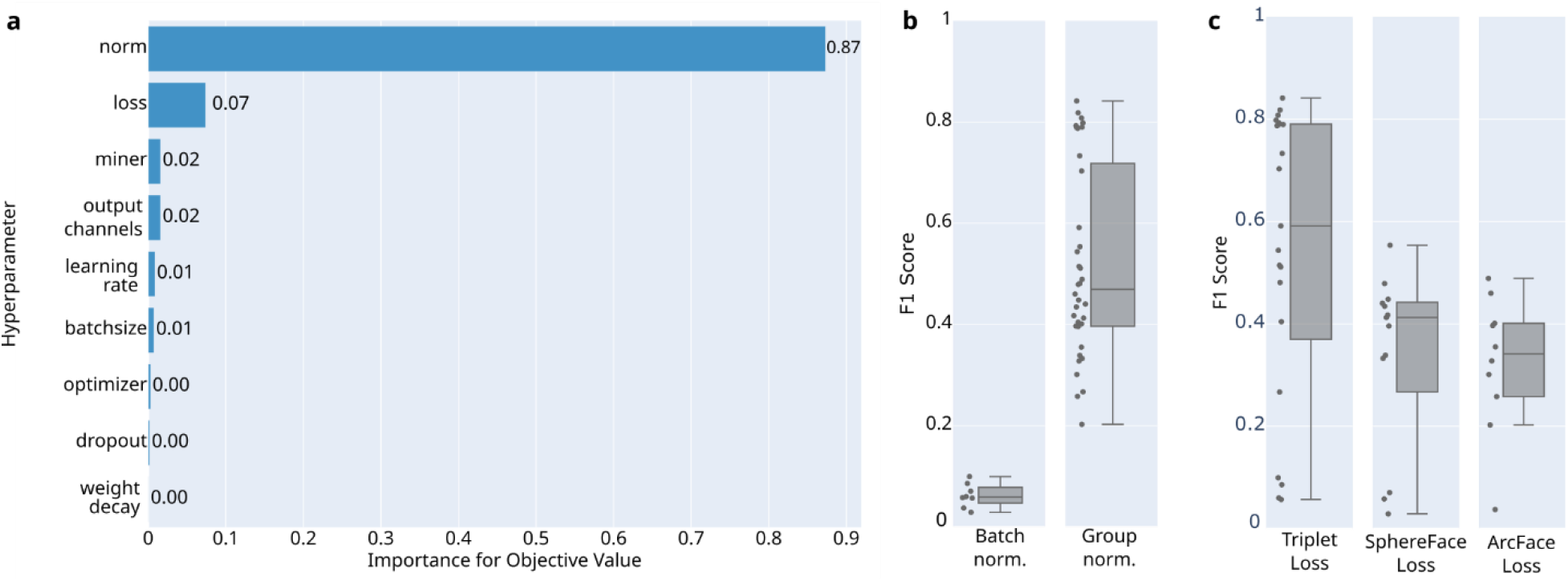
Hyperparameter optimization of TomoTwin. **a,** Hyperparameter importance estimated by Optuna^50^ after 180 trials with different configurations. **b,** F1 scores for trials using either the batch normalization or group normalization layers in convolutional neural network. Points represent the individual trials. Group normalization performed in general better than batch normalization in all cases. **c,** F1 score for trials using either Triplet-, SphereFace-, or ArcFace-Loss.

